# Cryo-EM structure of the human PAC1 receptor coupled to an engineered heterotrimeric G protein

**DOI:** 10.1101/2019.12.23.887737

**Authors:** Kazuhiro Kobayashi, Wataru Shihoya, Tomohiro Nishizawa, Francois Marie Ngako Kadji, Junken Aoki, Asuka Inoue, Osamu Nureki

**Affiliations:** Department of Biological Sciences, Graduate School of Science, The University of Tokyo, Bunkyo, Tokyo 113-0033, Japan; Graduate School of Pharmaceutical Sciences, Tohoku University, 6-3, Aoba, Aramaki, Aoba-ku, Sendai, Miyagi 980-8578, Japan

## Abstract

Pituitary adenylate cyclase-activating polypeptide (PACAP) is a pleiotropic neuropeptide hormone functioning in the central nervous system and peripheral tissues. The PACAP receptor PAC1R, which belongs to the class B G-protein-coupled receptors (GPCRs), is a drug target for mental disorders and dry eye syndrome. Here we present a cryo-electron microscopy structure of human PAC1R bound to PACAP and an engineered Gs heterotrimer. The structure revealed that TM1 plays an essential role in PACAP recognition. The ECD (extracellular domain) of PAC1R tilts by ~40° as compared to that of the glucagon-like peptide-1 receptor (GLP1R), and thus does not cover the peptide ligand. A functional analysis demonstrated that the PAC1R-ECD functions as an affinity trap and is not required for receptor activation, whereas the GLP1R-ECD plays an indispensable role in receptor activation, illuminating the functional diversity of the ECDs in the class B GPCRs. Our structural information will facilitate the design and improvement of better PAC1R agonists for clinical applications.

This article is a preprint version and has not been certified by peer review.

## Introduction

Pituitary adenylate cyclase-activating polypeptide (PACAP), a 38-amino acid linear peptide discovered in extracts of ovine hypothalamus^1^, is a multi-functional peptide hormone that acts as a neurotrophic factor, neuroprotectant, neurotransmitter, immunomodulator, and vasodilator^2^. PACAP is distributed mainly in the central nervous system (CNS), but is also detected in the testis, adrenal gland, digestive tract, and other peripheral organs. PACAP shares 68% amino acid sequence homology with vasoactive intestinal polypeptide (VIP). PACAP and VIP stimulate three different PACAP receptors: PAC1R^3^, VPAC1R, and VPAC2R, with different affinities. These receptors share about 50% sequence identity. The affinity of PAC1R for PACAP is higher than that for VIP^4^, indicating that PAC1R is relatively selective for PACAP.

PAC1R belongs to the class B G-protein-coupled receptors (GPCRs), and predominantly activates the adenylyl cyclase stimulatory G protein Gs. PAC1R is widely expressed in the CNS and peripheral tissues^2^. PACAP/PAC1R signaling has been implicated in playing essential roles in several cellular processes, including circadian rhythm regulation, food intake control, glucose metabolism, learning and memory, neuronal ontogenesis, apoptosis, and immune system regulation. Furthermore, perturbations in the PACAP/PAC1R pathway cause abnormal stress responses underlying posttraumatic stress disorder (PTSD)^5^, and thus PAC1R has been studied as a drug target for numerous disorders. PACAP and PAC1R are expressed in lacrimal glands, and induce tear secretion by increasing the aquaporin 5 (AQP5) levels in the plasma membrane^6^. Therefore, PAC1R is also a drug target for dry eye syndrome. However, the design of small molecule agonists for PAC1R has not yet been achieved, limiting the clinical applications targeting PAC1R.

PAC1R comprises two distinct domains: an N-terminal extracellular domain (ECD) and a transmembrane domain (TMD), as in the other class B GPCRs. A two-step/two-domain model has been proposed for ligand binding and receptor activation in the class B GPCRs^7^: the ECD is responsible for the initial and high-affinity binding of peptide ligands, and the TMD plays a key role in both ligand binding and receptor activation. A previous study suggested that PAC1R follows this model, and the PAC1R-ECD is not required for receptor activation^8^. However, in glucagon-like peptide 1 receptor (GLP1R), the ECD also plays an indispensable role in receptor activation, suggesting the divergent role of the ECD in the activation of class B GPCRs. Although the crystal structure of the PAC1R-ECD was determined in a ligand-free conformation^9^, little is known about the mechanism of the ligand recognition and signal transduction by PAC1R. Here we present a cryo-electron microscopy (Cryo-EM) structure of the human PAC1R, bound to the endogenous ligand PACAP and coupled to an engineered Gs heterotrimer. The structure, combined with complementary functional analyses, revealed the unique interaction between PACAP and the PAC1R-TMD and the structural basis for the functional divergence between the PAC1R-ECD and GLP1R-ECD.

## Results

### Overall structure

To facilitate expression and purification, we truncated the C-terminal residues 418–468 of the human PAC1R. This truncation did not alter the Gs-coupling activity, as measured by a NanoBiT-G-protein dissociation assay^10^ (Supplementary Fig. 1a, b, and Table 1). For the cryo-EM analysis, we used the mini-Gs protein, an engineered minimal G protein developed for structural studies^11^. The C-terminal truncated PAC1R was purified in the presence of PACAP and subsequently incubated with the mini-Gs heterotrimer (mini-Gs, β1, and γ2) and the nanobody Nb35, which stabilizes the GPCR-Gs complex. The reconstructed complex was purified by gel filtration.

**Table 1.** Pharmacological characterization of mutant PAC1Rs.

| | PAC1R-WT | ΔC | G389A | P360A | G363A |
| --- | --- | --- | --- | --- | --- |
| n = | 4 | 3 | 4 | 4 | 4 |
| EC<sub>50</sub> (nM) | 0.73 | 0.38 | 1.2 | 0.83 | > 1 μM |
| pEC<sub>50</sub> (mean ± SEM) | 9.14 ± 0.22 | 9.42 ± 0.03 | 8.92 ± 0.20 | 9.08 ± 0.19 | < 6 |

Vitrified complexes were imaged using a Titan Krios microscope equipped with a VPP (Supplementary Fig. 2). The 3D classification revealed two different classes, one containing a single complex (monomer class) and the other containing two complexes with inverted molecular packing (dimer class), which probably formed during the sample preparation. The structures of these two classes were determined at 4.5 Å and 4.0 Å resolutions, respectively, with the gold-standard Fourier shell correlation (FSC) criteria. Since the cryo-EM density suggested almost identical conformations in these classes, we built the atomic model of the receptor, ligand, and G-protein based on the higher resolution dimer class cryo-EM map. The local resolution of the map reached about 3.7 Å in the core region, including the TM helices of the receptor and the α5 helix of the Gαs Ras-like domain (Fig. 1a, b, Table 2, and Supplementary Fig. 3a). The molecular packing of the two complexes in the dimer class is solely mediated through a weak hydrophobic contact between V318^5.48^ and M322^5.52^ (Wooten numbering in superscript) in TM5, and the ECD and G-protein are not engaged in this interaction (Supplementary Fig. 3b), indicating that the dimerization minimally affects the conformation of the Gs-complexed PAC1R structure.

**Fig. 1.**
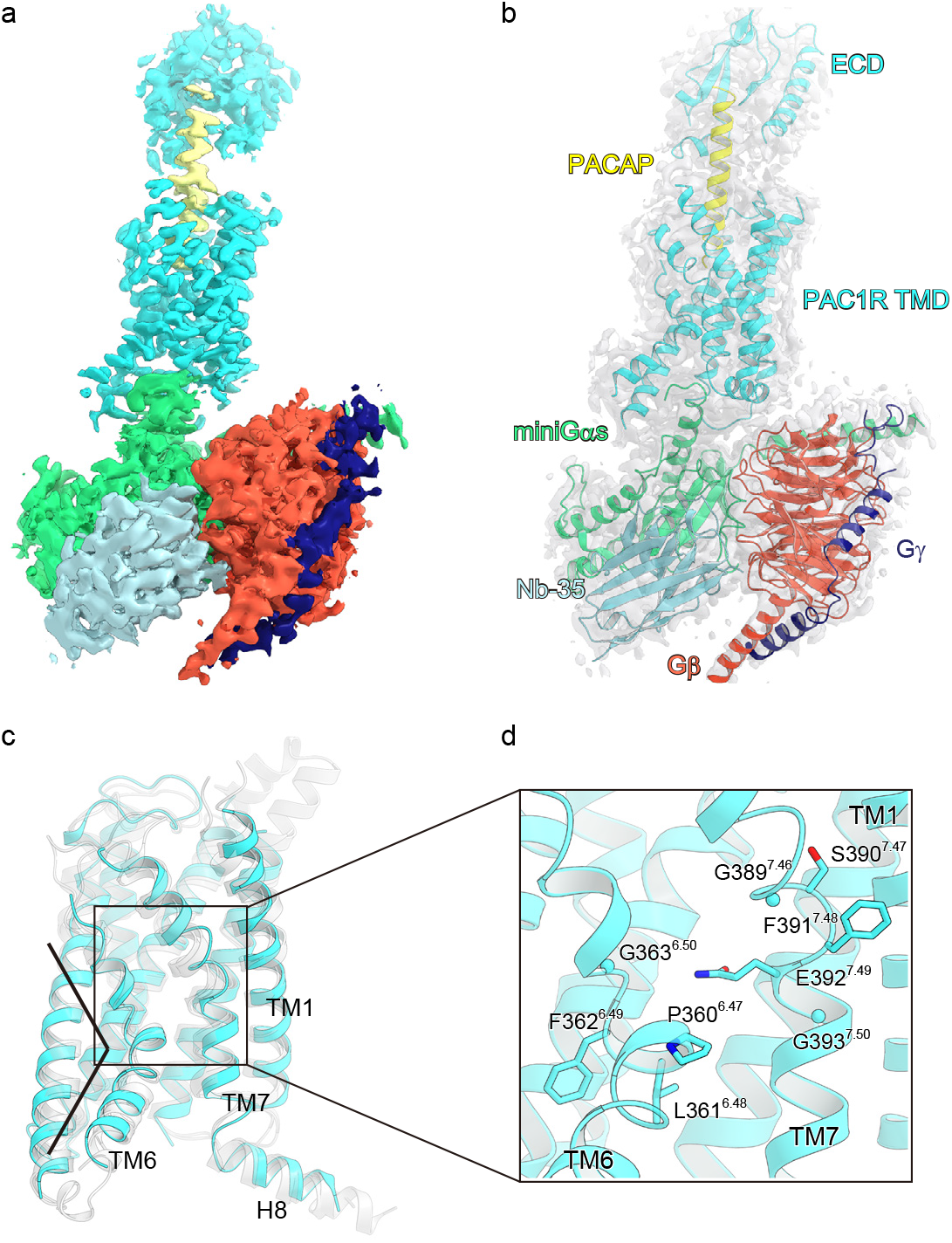
Overall structure of the PAC1R-mini-G_S_β_1_γ_2_-Nb35 complex. **a**, Sharpened cryo-EM map with variably colored densities (PAC1R TM: cyan, PACAP: yellow, mini-Gs heterotrimer: green, red, and purple, Nb35: light blue). **b**, Structure of the complex determined after refinement in the cryo-EM map. The model is shown as a ribbon representation with the transparent map. **c**, Superimposition of the TMD structures of PAC1R (cyan) and the other class B GPCRs determined to date (gray). **d**, TM7 unwinding in the PAC1R structure. Residues 387 to 393 are shown as sticks.

**Table 2.** Data collection, processing, model refinement, and validation.

|  |  |
| --- | --- |
| **Data collection and processing** | FEI Titan Krios |
| Magnification | 96,000 |
| Voltage (kV) | 300 |
| Electron exposure (e<sup>-</sup> Å<sup>-2</sup>) | 64 |
| Defocus range (μm) | -0.8 to -1.6 |
| Pixel size (Å)<sup>a</sup> | 0.861 |
| Symmetry imposed | C2 |
| Initial particle images (no.) | 980,964 |
| Final particle images (no.) | 132,808 |
| Map resolution (Å) | 4.0 |
| FSC threshold | 0.143 |
| Map resolution range (Å) | 3.7–4.9 |
| **Refinement** |  |
| Initial model used (PDB code) | 3N94, 6B3J |
| Model resolution (Å) | 4.0 |
| FSC threshold | 0.143 |
| Model resolution range (Å) |  |
| Map sharpening *B* factor (Å<sup>2</sup>) | -162.683 |
| **Model composition** |  |
| Non-hydrogen atoms | 8842 |
| Protein residues | 1081 |

| R.m.s. deviations |  |
| --- | --- |
| Bond lengths (Å) | 0.010 |
| Bond angles (°) | 1.162 |
| Validation |  |
| Clashscore | 6.46 |
| Rotamer outliers (%) | 1.88 |
| Ramachandran plot |  |
| Favored (%) | 92.14 |
| Allowed (%) | 7.86 |
| Disallowed (%) | 0 |

The PAC1R-TMD adopts the typical architecture of the activated class B GPCR conformation^12–16^, characterized by a sharp kink at TM6 (Fig. 1c). One notable difference is observed in TM7, which is kinked at the highly conserved G393^7.50^ in the other class B GPCRs. PAC1R has an additional glycine G389^7.46^ near G393^7.50^, and thus TM7 unwinds and bends around G389^7.46^ in the current structure (Fig. 1d). However, the G389^7.46^A mutation, which would facilitate the α-helical formation of the unwound TM7, did not alter the Gs-coupling activity (Supplementary Fig. 1a, b, and Table 1). This result suggests that this unwinding in TM7 is not related to the PAC1R function.

### Interaction between PACAP and PAC1R-TMD

We observed an unambiguous density extending from the TMD, which allowed us to assign the secondary structure and side-chain orientations of PACAP (Fig. 2a). The N-terminus of the peptide ligand PACAP is directed toward the TMD core, as in the other class B GPCR structures. The H1 to L27 residues of PACAP form a continuous α-helix and protrude from the transmembrane binding pocket. By contrast, the residues after G28 are disordered, consistent with the fact that the C-terminal truncated variant PACAP_1-27_ has the same affinity as PACAP. Notably, the N-terminal four residues (H1 to G4) form a continuous α-helix, together with the I5 to L27 residues, while these residues were disordered in the previous nuclear magnetic resonance (NMR) structure of PACAP_1-27_ bound to detergent micelles^17^. These residues are recognized by 13 residues of the receptor, and G4 of PACAP closely contacts the W306^5.36^ side chain of the receptor (Fig. 2b, c). These interactions stabilize the α-helical structure at the N-terminus of PACAP, which is essential for receptor activation.

**Fig. 2.**
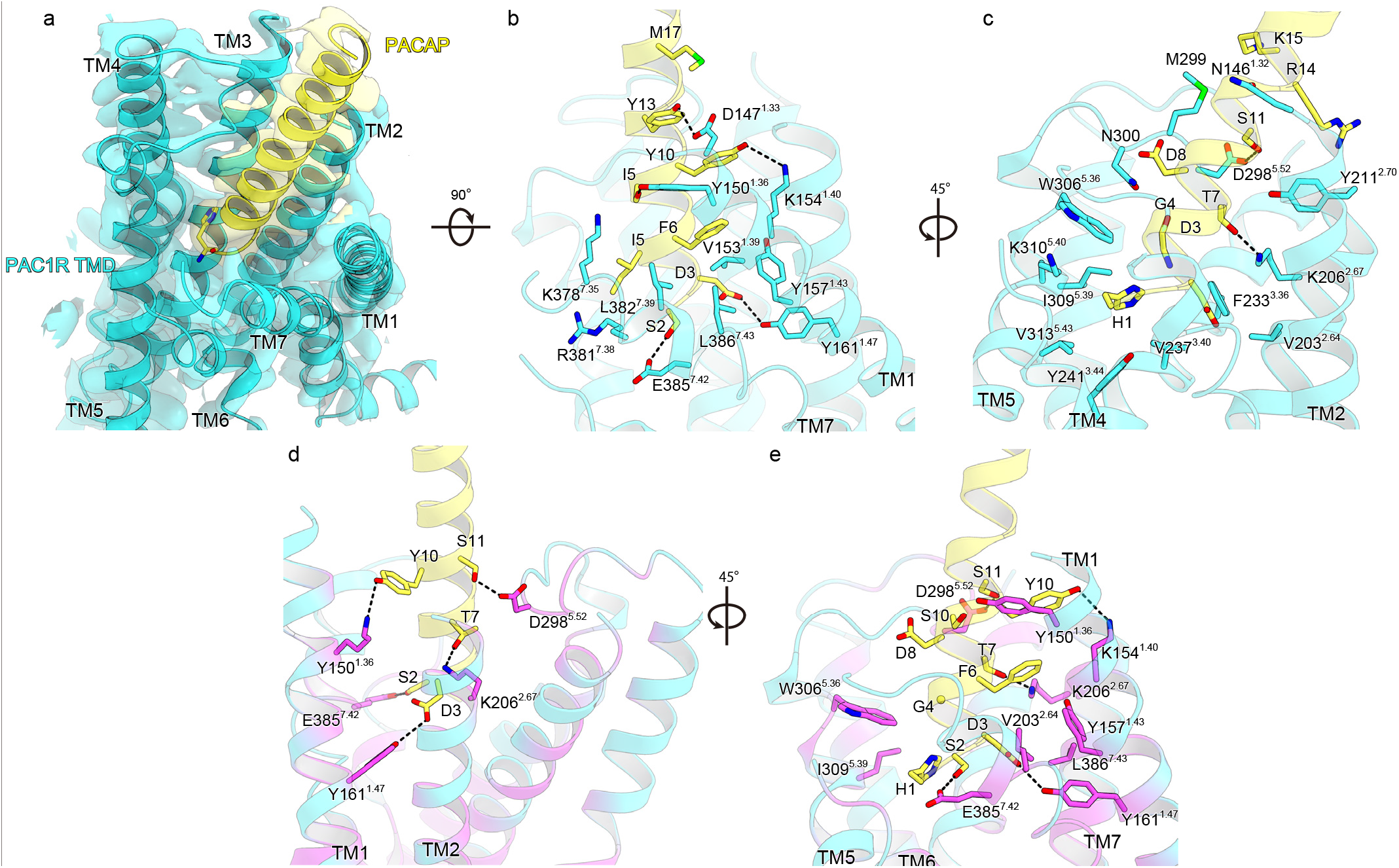
PACAP binding site in TMD. **a**, Sharpened map of PACAP and the TMD, viewed from the extracellular side. PACAP and TMD are shown as ribbon representations with the transparent map. **b, c**, Detailed interactions between PACAP and the TMD, shown as ribbon representations colored as in Fig. 1. Contact residues are shown as sticks. The interactions with TM1, 6, and 7 are shown in (**b**), while those with TM2, 3, and 5 are shown in (**c**). Hydrogen-bonding interactions are indicated by black dashed lines. **d, e**, Sequence conservation of the PACAP binding site between three types of PACAP receptors (PAC1R, VPAC1R, and VPAC2R), mapped onto the PAC1R structure. Conserved and non-conserved residues are colored magenta and cyan, respectively. The conserved hydrogen-bonding interactions are shown in (**d**), and all of the conserved residues are shown in (**e**).

The N-terminal 17 residues of PACAP create an extensive interaction network with TM 1-3, 5, 7, and ECL2 of the receptor (Fig. 2b, c). The details are summarized in Supplementary Table 1. Notably, PACAP forms numerous interactions with the extracellular portion of TM1, involving the aromatic residues of PACAP (F6, Y10, and Y13), PAC1R (Y150^1.36^, Y157^1.43^, and Y161^1.47^), and four hydrogen-bonding interactions (D3-Y161^1.47^, S9-Y150^1.36^, Y10-K154^1.40^, and Y13-D147^1.33^) (Fig. 2b-e). These close interactions with TM1 are not observed in the other class B GPCR structures (Supplementary Fig. 4a-d), and are a unique feature of the PACAP-PAC1R structure.

PACAP can activate three types of PACAP receptors, PAC1R, VPAC1R, and VPAC2R with similar affinities^4^. To investigate the similarity in their ligand recognition, we mapped the conserved residues on the current structure (Fig. 2d, e, and Supplementary Fig. 5). Notably, the residues involved in the ligand recognition are highly conserved in TM1, suggesting that TM1 plays a critical role in the PACAP recognition by the receptors.

## Structural insight into G-protein activation

In the class B GPCRs, ligand binding induces the rearrangement of the central polar interaction network, followed by the unwinding of TM6 at the highly conserved P^6.47^-X-X-G^6.50^ motif and the opening of the intracellular cavity of the receptor for G-protein coupling^12,13,14^. In the central region of PAC1R, we observed a similar polar interaction network and the unwinding of TM6 (Fig. 3a, b). The polar interaction network comprises D3 of PACAP and Y161^1.47^, R199^2.60^, N240^3.43^, Y241^3.44^, P360^6.47^, G363^6.50^, H365^6.52^, Y366^6.53^, and Q392^7.49^ of the receptor. Notably, D3 forms a hydrogen bond with Y161^1.47^ and an electrostatic interaction with R260^2.60^. R260^2.60^ in turn forms a hydrogen bond with Y241^3.44^. Y366^6.53^, in the extracellular portion of TM6, is directed toward the receptor core and participates in this network. Overall, this polar interaction network extends from D3 to the carbonyl oxygens of P360^6.47^ and G363^6.50^ in TM6. A previous SAR study showed that the substitution of D3 with alanine reduces both the ^E^max value to 70% and the affinity for the receptor^4^. Therefore, PACAP binding directly induces the rearrangement of the polar interaction network in the central region and plays a key role in receptor activation, by unwinding TM6.

**Fig. 3.**
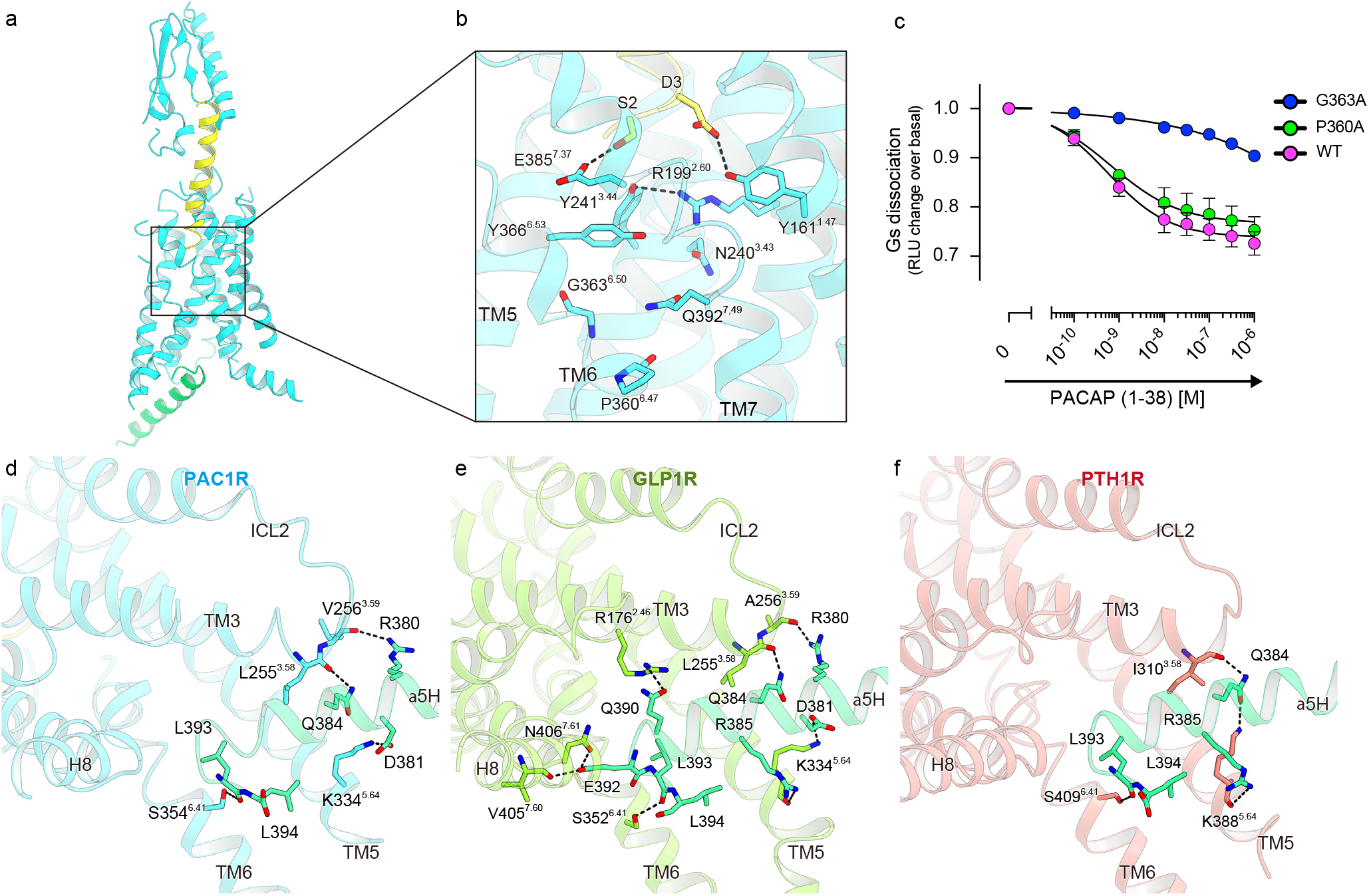
Mechanism of receptor activation and Gs coupling. **a**, Ribbon representation of the PAC1R-TMD, PACAP, and α5-helix of mini-Gs, viewed from the membrane plane and colored as in Fig. 1. **b**, The residues involved in the central polar interaction network are represented by sticks. Hydrogen bonding interactions are indicated by black dashed lines. **c**, PAC1R-mediated Gs activation, measured by the NanoBiT-G-protein dissociation assay. Cells transiently expressing the NanoBiT-Gs along with the indicated PAC1R construct were treated with PACAP (1-38), and the change in the luminescent signal was measured. **d-f**, Cytoplasmic views of the PAC1R (cyan) with the C-terminal α5 helix of Gαs (yellow) (**d**), compared to the GLP1-GLP1R:Gs complex (orange, PDB 5VAI) (**e**) and the LA-PTH-PTH1R:Gs complex (pink, PDB 6NBF) (**f**). Hydrogen bonding interactions are indicated by black dashed lines.

TM6 is kinked at P360^6.47^ and G363^6.50^ in the P^6.50^-X-X-G^6.53^ motif, as in the other Gs-complexed class B GPCR structures. Notably, the kink at G363^6.50^ is sharp (~ 90°), whereas that at P360^6.47^ is to a less extent (Fig. 3b). Previous mutational studies of the calcitonin receptor family members suggested the functional importance of P^6.47^ in receptor activation^18,19^. However, the P360^6.47^A mutation to PAC1R did not alter the Gs-coupling activity (Fig. 3c and Table 1). By contrast, the G363^6.50^A mutation completely abolished the activity, suggesting that G363^6.50^, rather than P360^6.47^, is responsible for the TM6 unwinding upon receptor activation, consistent with the structural observations.

The intracellular cavity of the receptor closely contacts the α5-helix of Gs, which is the primary determinant for the G-protein coupling (Fig. 3a, d). Specifically, S354^6.41^ in TM6 directly hydrogen bonds with the carbonyl oxygen of L393. K334^5.64^ in TM5 forms a salt bridge with D381, and the carbonyl oxygens of L255 and V256 in ICL2 hydrogen bond with Q384 and K380, respectively. These interactions are also observed in other Gs-complexed class B GPCR structures^13,16^ (Fig. 3e, f), suggesting that they are conserved structural features of the Gs-coupling in class B GPCRs.

### Diverged functional role of ECDs in class B GPCRs

The class B GPCRs have an ECD (about 120 amino acids) at the N-terminus, which is commonly important for the initial, high-affinity binding to peptide hormones. Although the ECD is less well resolved in our EM map, probably due to its flexibility, we could fit the ECD region of the previous PAC1R-ECD crystal structure (PDB code: 3N94)^9^ onto the map by a rigid body. This model can facilitate discussions about the interactions between PACAP and the ECD (Fig. 4a). The PAC1R-ECD adopts a three-layer α–β–βα fold, which is conserved in the class B GPCRs. The C-terminal portion of PACAP (Q16, V19, Y22, L23, and L27) interacts with the loops connecting β1-β2 and β3-β4, and the N-terminal ends of α-helix 1 and α-helix 2 in the PAC1R-ECD, as in other class B GPCRs (Supplementary Fig. 4e-g). Furthermore, the PAC1R-ECD has an additional α-helix, α-helix 3, in the loop connecting β3-β4, which closely contacts PACAP. A previous study showed that the N-terminal splice variant PAC1R-short^20^, which lacks residues 89-109 between the α-helix 3 and β4, exhibits increased affinity for PACAP. While residues 89-110 are not modeled in our EM map, we suggest that the truncation affects the conformation of the α-helix 3 and enhances the interaction with PACAP.

**Fig. 4.**
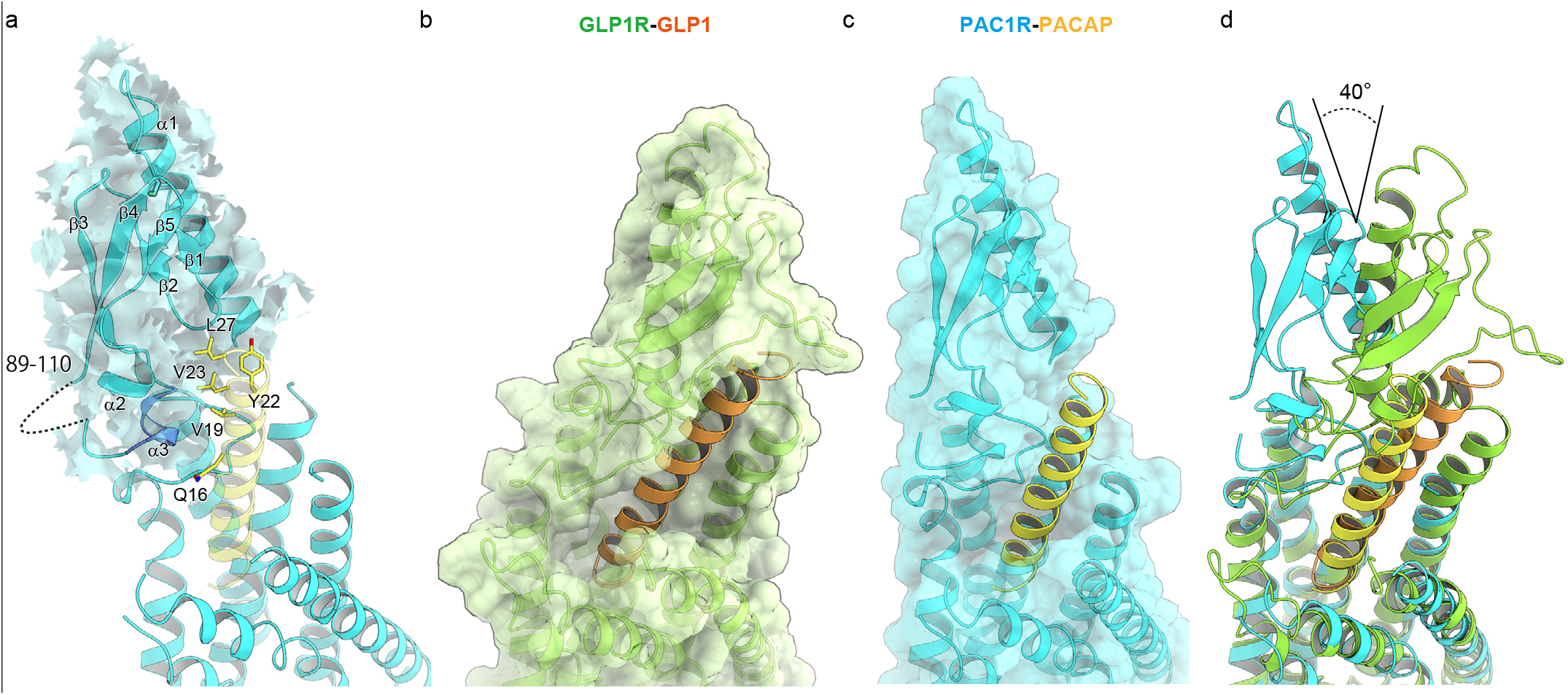
Structural comparison of PAC1R and GLP1R. **a**, Interaction between PACAP and the PAC1R-ECD. The PAC1R-ECD is shown as a ribbon representation with the transparent sharpened map. C54 is shown as a stick model. **b, c**, Surface representations of GLP1R (**b**) and PAC1R (**c**). The receptors are shown as ribbon representations with transparent surfaces. PACAP and PAC1R are colored as in Fig. 1a. GLP1 and GLP1R are colored orange and light-green, respectively. **d**, Superimposition of PAC1R and GLP1R, viewed from the membrane plane.

The ECD in class B GPCRs also plays a key role in receptor activation. Previous functional analyses demonstrated that the ECD-truncated GLP1R does not respond to GLP1^21^. In the GLP1R structure, the ECD covers the top of the C-terminal portion of GLP1, to facilitate the interactions between the N-terminal portion of GLP1 and TMD (Fig. 4b). However, the PAC1R-ECD tilts by ~40° as compared with the GLP1R-ECD (Fig. 4c, d), and thus it only interacts with the side of the C-terminal portion of PACAP, suggesting the different role of the PAC1R-ECD.

To investigate the function of the PAC1R-ECD, we truncated the C-terminal portion of PACAP (residues 18-38) that interacts with the ECD (Fig. 5a). The truncated peptide PACAP_1-17_ activated the receptor at the same level, as compared with PACAP in the NanoBiT-G-protein dissociation assay, while its EC_50_ was significantly increased by about 6000-fold (Fig. 5b and Table 3), suggesting that PACAP_1-17_ is capable of functioning as a full agonist for PAC1R. Moreover, PACAP and PACAP_1-17_ also activated the ECD-truncated PAC1R to mostly the same level (Fig. 5c, Table 3, and Supplementary Fig. 1b). These results indicate that the PAC1R-ECD functions merely as an affinity trap to bind and precisely localize the peptide hormone to the receptor, whereas the interaction between PACAP and the PAC1R-TMD is necessary and sufficient for receptor activation. This observation is consistent with the previous study, which showed that the PAC1R-TMD covalently linked to the PACAP_1-12_ at the N-terminus constitutively activates the G-protein21. By contrast, GLP1_7-23_, which lacks the C-terminal portion of GLP (Fig. 5d), completely lost the agonist activity for GLP1R (Fig. 5e and Table 3). Furthermore, the ECD-truncated GLP1R was poorly expressed and lacked receptor activity (Fig. 5f and Supplementary Fig. 1c). These results confirmed that the GLP1R-ECD plays an indispensable role in receptor activation. While the ECDs are commonly essential for ligand recognition in the class B GPCRs, their contributions to receptor activation diverge among the receptors.

**Fig. 5.**
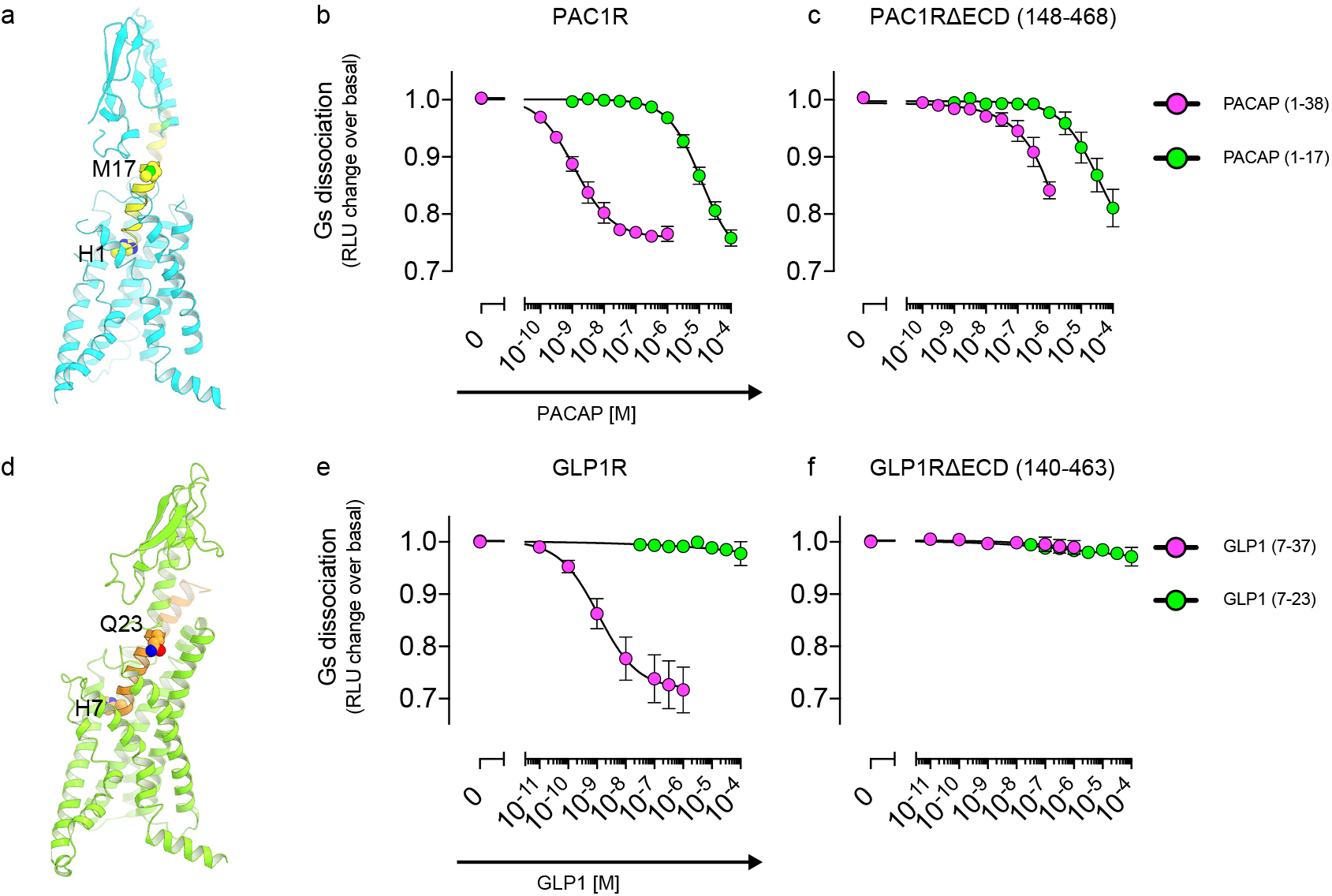
Characterization of the truncated analogs of PACAP and GLP1. **a**, Overall structure of the PACAP-bound PAC1R, viewed from the membrane plane. PACAP and PAC1R are shown as ribbon representations, colored as in Fig. 1a. The C-terminal portion of PACAP (18-38) is shown with increased transparency. **b, c**, PACAP-induced Gs activation measured by the NanoBiT-G-protein dissociation assay. Cells transiently expressing the NanoBiT-Gs along with PAC1R (b) or PAC1R∆ECD (c) were stimulated by the indicated PACAP peptides, and the change in the luminescent signal was measured. **d**, Overall structure of the GLP1-bound GLP1R, viewed from the membrane plane. GLP1 and GLP1R are shown as ribbon representations, colored as in Fig. 4b. The C-terminal portion of GLP1 (24-37) is shown with increased transparency. **e, f**, GLP-1-induced Gs activation measured by the NanoBiT-G-protein dissociation assay. Cells transiently expressing the NanoBiT-Gs along with GLP1R (e) or GLP1R∆ECD (f) were stimulated by the indicated GLP-1 peptides, and the change in the luminescent signal was measured.

**Table 3.** Pharmacological characterization of truncated analogs of PACAP and GLP1

|  |  | PAC1R-WT | PAC1R-TMD |
| --- | --- | --- | --- |
|  | n = | 6 | 4 |
| PACAP<sub>1-38</sub> | EC<sub>50</sub> (nM) | 1.3 | 580 |
|  | pEC<sub>50</sub> (mean ± SEM) | 8.90 ± 0.10 | 6.24 ± 0.08 |
| PACAP<sub>1-17</sub> | EC<sub>50</sub> (μM) | 8.1 | 19 |
|  | pEC<sub>50</sub>, (mean ± SEM) | 5.09 ± 0.09 | 4.72 ± 0.17 |
|  |  | GLP1R-WT | GLP1R-TMD |
|  | n = | 4 | 3 |
| GLP1<sub>7-37</sub> | EC<sub>50</sub> (nM) | 1.2 | >1000 |
|  | pEC<sub>50</sub> (mean ± SEM) | 8.93 ± 0.17 | < 6 |
| GLP1<sub>7-23</sub> | EC<sub>50</sub> (μM) | >100 | >100 |
|  | pEC<sub>50</sub>, (mean ± SEM) | <4 | <4 |

## Discussion

We determined the PAC1R structure in complex with PACAP and the Gs-protein, which revealed a unique interaction between PACAP and the PAC1R-TMD, involving the aromatic residues in PACAP and TM1. Structural observations and functional analyses indicated that the interaction between PACAP and the TMD is necessary and sufficient for receptor activation, while the ECD is only required for the high-affinity binding. Our structural information will help the design of novel peptide-mimetic agonists for PAC1R, to treat dry eye syndrome and mental disorders.

The class B GPCRs include 15 receptors in humans and are commonly activated by peptide ligands. The class B receptors share similar characteristics, such as the N-terminal ECD, and are distinct from the class A GPCRs, which are activated by diverse ligands (e.g., peptides, amines, purines, and lipids)^22^. The structures of the class B GPCRs suggested two different types of ligand recognition. In the structures of calcitonin receptor^12^ and calcitonin receptor-like receptor (CLR)^15^, the N-terminal portions of the peptide ligands bind to the TMD in α-helical conformations, while the C-terminal portion binds to the ECD in an extended conformation (Supplementary Fig. 4d, h). The function of the calcitonin receptor family is modified by receptor activity-modifying proteins (RAMPs). Essentially, CLR can receive calcitonin gene-related peptide (CGRP) by the interaction between its ECD and RAMP1, suggesting that the ECD plays a key role in both ligand binding and receptor activation. In the structures of the glucagon receptor family members, GLP1R^13,14^ and glucagon receptor, GCGR^23,24^, the peptide ligands adopt continuous α-helices and their ECDs cover the ligands to facilitate the interactions with the TMDs, thus playing an indispensable role in receptor activation. Although PACAP also adopts a continuous α-helix, the PAC1R-ECD has no functional role in receptor activation, because PACAP can interact with the TMD without the aid of the ECD. The PAC1R-ECD functions merely as an affinity trap for the high-affinity binding of PACAP. Despite the structural similarities in the class B GPCRs, the functional roles of the ECD are diverse.

## Acknowledgements

We thank R. Danev and M. Kikkawa for setting up the cryo-EM infrastructure, K. Ogomori for technical assistance, and K. Yamashita for model building. We also thank Ayumi Inoue (Tohoku University, Japan) for technical assistance. This work was supported by grants from the Platform for Drug Discovery, Informatics and Structural Life Science by the Ministry of Education, Culture, Sports, Science and Technology (MEXT), JSPS KAKENHI grants 16H06294 (O.N.), 17J30010 (W.S.), 30809421 (W.S.), 17K08264 (A.I.), 17H05000 (T.N.) and the Japan Agency for Medical Research and Development (AMED) grants: the PRIME JP18gm5910013 (A.I.) and the LEAP JP18gm0010004 (A.I. and J.A.), and the National Institute of Biomedical Innovation.

## Author contributions

K.K. expressed and purified the mini-Gs heterotrimer, and performed the complex formation, grid-preparation, and cryo-EM observation. W.S. designed the experiments, purified the receptor, established the preparation method for the mini-Gs heterotrimer and Nb35, and refined the structure. T.N. performed the cryo-EM data collection and single particle analysis. A.I., F.M.N.K., and J.A. performed and oversaw the cell-based assays. The manuscript was mainly prepared by W.S., K.K., and A.I., with assistance from T.N. and O.N.

## Expression and purification of the human PAC1R

The N-terminal signal sequence in human PAC1R (Genbank ID: AK290046) was replaced with the haemagglutinin signal peptide. The C-terminus was truncated after S417. The modified receptor was subcloned into a modified pFastBac vector^25^, with the resulting construct encoding a TEV cleavage site followed by a GFP-His^10^ tag at the C-terminus. The recombinant baculovirus was prepared using the Bac-to-Bac baculovirus expression system (Invitrogen). Sf9 insect cells were infected with the virus at a cell density of 4.0 × 10^6^ cells per milliliter in Sf900 II medium, and grown for 48 h at 27 °C. The harvested cells were disrupted by sonication, in buffer containing 20 mM Tris-HCl, pH 7.5, and 20% glycerol. The crude membrane fraction was collected by ultracentrifugation at 180,000*g* for 1 h. The membrane fraction was solubilized in buffer, containing 20 mM Tris-HCl, pH 7.5, 200 mM NaCl, 1% LMNG, 0.1 %CHS, 20% glycerol, and 1 μM PACAP38, for 2 h at 4 °C. The supernatant was separated from the insoluble material by ultracentrifugation at 180,000 *g* for 20 min, and incubated with TALON resin (Clontech) for 30 min. The resin was washed with ten column volumes of buffer, containing 20 mM Tris-HCl, pH 7.5, 500 mM NaCl, 0.01% LMNG, 0.001% CHS, 0.1 μM PACAP38, and 15 mM imidazole. The receptor was eluted in buffer, containing 20 mM Tris-HCl, pH 7.5, 500 mM NaCl, 0.01% LMNG, 0.001% CHS, 0.1 μM PACAP38, and 200 mM imidazole. The eluate was treated with TEV protease and dialyzed against buffer (20 mM Tris-HCl, pH 7.5, 500 mM NaCl). The cleaved GFP–His_10_ tag and the TEV protease were removed with Co^2+^-NTA resin. The receptor was concentrated and loaded onto a Superdex200 10/300 Increase size-exclusion column, equilibrated in buffer containing 20 mM Tris-HCl, pH 7.5, 150 mM NaCl, 0.01% LMNG, 0.001% CHS, and 0.1 μM PACAP38. Peak fractions were pooled, concentrated to 5 mg ml^−1^ using a centrifugal filter device (Millipore 50 kDa MW cutoff), and frozen in liquid nitrogen.

### Expression and purification of the mini-Gs heterotrimer

The gene encoding mini-Gs^26^, with codons optimized for an *E. coli* expression system, was synthesized (GeneArt) and subcloned into a modified pET21a(+)-vector, with the resulting construct encoding a His_6_ tag followed by a TEV cleavage site at the N-terminus. The protein was expressed in *E. coli* BL21 cells. Protein expression was induced by 1 mM isopropyl β-D-thiogalactopyranoside (IPTG) for 20 h at 25 °C. The harvested cells were disrupted by sonication, in buffer containing 20 mM Tris-HCl, pH 7.5, 20% glycerol, 10 μM GDP, and 10 mM imidazole. The cell debris was removed by centrifugation at 25,000*g* for 30 min. The supernatant was incubated with Ni-NTA resin (Qiagen) for 30 min. The resin was washed with ten column volumes of buffer, containing 20 mM Tris-HCl, pH 7.5, 500 mM NaCl, 10 μM GDP, and 30 mM imidazole. The protein was eluted in buffer, containing 20 mM Tris-HCl, pH 7.5, 500 mM NaCl, 10 μM GDP, and 200 mM imidazole. The eluate was treated with TEV protease and dialyzed against buffer (20 mM Tris-HCl, pH 7.5, 150 mM NaCl, and 10 μM GDP). The TEV protease was removed by Ni-NTA resin. The protein was concentrated and loaded onto a Hiload Superdex200 10/300 Increase size-exclusion column, equilibrated in buffer containing 20 mM Tris-HCl, pH 7.5, 150 mM NaCl, and 1 μM GDP). Peak fractions were pooled, concentrated to 8 mg ml^−1^ using a centrifugal filter device (Millipore 10 kDa MW cutoff), and frozen in liquid nitrogen.

His_6_-rat Gβ_1_ and bovine Gγ_2_ were subcloned into the pFastBac Dual vector. The recombinant baculovirus was prepared using the Bac-to-Bac baculovirus expression system (Invitrogen). Sf9 insect cells were infected with the virus at a cell density of 4.0 × 10^6^ cells per milliliter in Sf900 II medium, and grown for 48 h at 27 °C. The harvested cells were disrupted by sonication, in buffer containing 20 mM Tris-HCl, pH 7.5, 150 mM NaCl, 10 mM imidazole, and 20% glycerol, and clarified by ultracentrifugation at 180,000*g* for 30 min. The supernatant was incubated with Ni-NTA resin (Qiagen) for 30 min. The resin was washed with ten column volumes of buffer, containing 20 mM Tris-HCl, pH 7.5, 500 mM NaCl, and 30 mM imidazole. The protein was eluted in buffer, containing 20 mM Tris-HCl, pH 7.5, 500 mM NaCl, and 200 mM imidazole. The protein was concentrated and loaded onto a Superdex200 10/300 Increase size-exclusion column, equilibrated in buffer containing 20 mM Tris-HCl, pH 7.5, and 150 mM NaCl. Peak fractions were pooled, concentrated to 8 mg ml^−1^ using a centrifugal filter device (Millipore 10 kDa MW cutoff), and frozen in liquid nitrogen.

The purified mini-Gs and Gβ_1_Gγ_2_ were mixed and incubated overnight on ice. The sample was concentrated and loaded onto a Superdex200 10/300 Increase size-exclusion column, equilibrated in buffer containing 20 mM Tris-HCl, pH 7.5, 150 mM NaCl, and 1 μM GDP. The fractions containing the mini-Gs heterotrimer were pooled, concentrated to 8 mg ml^−1^ using a centrifugal filter device (Millipore 10 kDa MW cutoff), and frozen in liquid nitrogen.

## Expression and purification of Nb35

The gene encoding the C-terminally His_6_-tagged nanobody-35 (Nb35), with codons optimized for an *E. coli* expression system, was synthesized (GeneArt) and subcloned into the pET22b(+)-vector. The protein was expressed in the periplasm of *E. coli* C41(Rosetta) cells. Protein expression was induced by 1 mM isopropyl β-D-thiogalactopyranoside (IPTG) for 20 h at 25 °C. The harvested cells were disrupted by sonication, in buffer containing 20 mM Tris-HCl, pH 7.5, and 20% glycerol. The cell debris was removed by centrifugation at 25,000*g* for 30 min. The supernatant was incubated with Ni-NTA resin (Qiagen) for 30 min. The resin was washed with ten column volumes of buffer, containing 20 mM Tris-HCl, pH 7.5, 500 mM NaCl, and 30 mM imidazole. The protein was eluted in buffer, containing 20 mM Tris-HCl, pH 7.5, 500 mM NaCl, and 200 mM imidazole. The eluate was dialyzed against buffer (20 mM Tris-HCl, pH 7.5, 150 mM NaCl). The protein was concentrated to 3 mg ml^−1^ using a centrifugal filter device (Millipore 10 kDa MW cutoff), and frozen in liquid nitrogen.

### Formation and purification of the PAC1R-mini-G_S_β_1_γ_2_-Nb35 complex

Purified PAC1R was mixed with a 1.2-fold molar excess of mini-Gsβ1γ2 and a 1.5-fold molar excess of Nb35 in the presence of apyrase (0.1 U/ml) and the mixture was incubated on ice overnight. The sample was loaded onto a Superdex200 10/300 Increase size-exclusion column, equilibrated in buffer containing 20 mM HEPES-Na, pH 7.5, 150 mM NaCl, 0.0075% LMNG, 0.0025% GDN, and 0.00025%CHS. Peak fractions of the PAC1R-mini-Gsβ_1_γ_2_-Nb35 complex were pooled and concentrated to 8 mg/ml.

## Sample vitrification and cryo-EM data acquisition

The purified complex was applied onto a freshly glow-discharged Quantifoil holey carbon grid (R1.2/1.3, Cu/Rh, 300 mesh), blotted for 4 s at 4 °C in 100% humidity, and plunge-frozen in liquid ethane by using a Vitrobot Mark IV. The grid images were obtained with a 300kV Titan Krios G3i microscope (Thermo Fisher Scientific), equipped with a GIF Quantum energy filter (Gatan), a Volta phase plate (Thermo Fisher Scientific), and a Falcon III direct electron detector (Thermo Fisher Scientific). A total of 2,895 movies were obtained in the electron counting mode, with a physical pixel size of 0.861 Å. The data set was acquired with the EPU software, with a defocus range of −0.8 to −1.6 μm. Each image was dose-fractionated to 64 frames at a dose rate of 6– 8 e^−^ pixel^−1^ per second, to accumulate a total dose of 64 e^−^ Å^−2^. In total, 2,895 super-resolution movies were collected.

## Image processing

The movie frames were aligned in 5 × 5 patches, dose weighted, and binned by 2 in MotionCor2^27^. Defocus parameters were estimated by CTFFIND 4.1^28^. First, template-based auto-picking was performed with the two-dimensional class averages of a few hundred manually picked particles as templates. A total of 980,964 particles were extracted in 3.24 Å pixel^−1^. These particles were subjected to three rounds of two-dimensional classification in RELION 3.0. The initial model was generated in RELION-3.0^28^. Subsequently, 980,964 particles were further classified in 3D without symmetry. Two stable classes showed detailed features for all subunits. One contained a single complex (monomer class). The other contained two complexes in an inverted molecular packing with C2 symmetry (dimer class). The particles of the monomer and dimer classes were 282,622 and 132,808 particles, respectively, were then re-extracted with the original pixel size of 1.35 Å pixel, and subsequently subjected to 3D refinement. The resulting 3D models and particle sets were subjected to per-particle defocus refinement, Bayesian polishing, and 3D refinement. The final 3D refinement and postprocessing yielded maps of the monomer and dimer classes with global resolutions of 4.5 Å and 4.0 Å, respectively. The comparison of the density maps of the two classes suggested almost identical conformation, and therefore, we built an atomic model onto the higher resolution map of the dimer class. All density maps were sharpened by applying the temperature-factor, which was estimated using the post-processing in RELION-3.1. The local resolution was estimated by RELION-3.1. The processing strategy is described in Supplementary figure 2.

## Model building and refinement

The initial template for the PAC1R transmembrane regions, PACAP, G-protein, and Nb35 was derived from the structure of human GLP1R in complex with a dominant-negative Gαs (PDB code: 6B3J), followed by extensive remodeling using COOT^29^. Owing to the discontinuous and/or variable density in the ECD region, we assigned the high-resolution X-ray crystal structure of the PAC1R (PDB code: 3N94)^9^ by a rigid body fit, and the model was rebuilt using Rosetta^30^ against the density, manually readjusted using COOT, and refined using phenix.real_space_refine^31^. Validation was performed in MolProbity^32^. The potential overfitting of the refined models was tested by using a cross-validation method, as described previously. Briefly, the final models were ‘shaken’ by introducing random shifts to the atomic coordinates with an rms of 0.5 Å, and were refined against the first half map. These shaken refined models were used to calculate the FSC against the same first half maps (FSC_half1_ or work), and the second half maps (FSC_half2_ or free) that were not used for the refinement, using phenix.mtriage. The small differences between the FSC_half1_ and FSC_half2_ curves indicated no severe overfitting of the models. The curves representing model vs. full map were calculated, based on the final model and the full, filtered and sharpened map. The statistics of the 3D reconstruction and model refinement are summarized in Table 2. All molecular graphics figures were prepared with CueMol (http://www.cuemol.org) and UCSF Chimera^33^.

## NanoBit G-protein dissociation assay

PAC1R- and GLP1R-induced Gs activation was measured by a NanoBiT-G-protein dissociation assay^10^, in which the interaction between a Gα subunit and a Gβγ subunit was monitored by a NanoBiT system (Promega). Specifically, a NanoBiT-Gs protein consisting of a large fragment (LgBiT)-containing Gαs subunit and a small fragment (SmBiT)-fused Gγ2 subunit, along with the untagged Gβ_1_ subunit, was expressed with a test GPCR, and the ligand-induced luminescent signal change was measured. We used the N-terminal FLAG (DYKDDDK) tagged constructs of the human PAC1R, PAC1RΔECD (148-468), GLP1R, and GLP1RΔECD (140-463). HEK293 cells deficient for G_q/11_34 were seeded in a 6-well culture plate at a concentration of 2 × 10^5^ cells ml^−1^ (2 ml per well in DMEM (Nissui Pharmaceutical) supplemented with 10% fetal bovine serum (Gibco), glutamine, penicillin, and streptomycin), one day before transfection. The transfection solution was prepared by combining 4 µl (per well hereafter) of polyethylenimine solution (Polysciences, 1 mg ml^−1^) and a plasmid mixture consisting of 100 ng LgBiT-containing Gαs subunit, 500 ng Gβ_1_, 500 ng SmBiT-fused Gγ_2_, and 200 ng test GPCR (or an empty plasmid) in 200 µl of Opti-MEM (ThermoFisher Scientific). To prepare a larger volume of transfected cells, 10-cm culture dishes (10 ml culture volume) were used with 5-fold scaling of the 6-well plate contents. After an incubation for one day, the transfected cells were harvested with 0.5 mM EDTA-containing Dulbecco’s PBS, centrifuged, and suspended in 2 ml of HBSS containing 0.01% bovine serum albumin (BSA fatty acid–free grade, SERVA) and 5 mM HEPES (pH 7.4) (assay buffer). The cell suspension was dispensed in a white 96-well plate at a volume of 80 µl per well, and loaded with 20 µl of 50 µM coelenterazine (Carbosynth), diluted in the assay buffer. After 1 h incubation at room temperature, the titrated antagonist (Atropine, NMS, or Tiotropium), diluted in the assay buffer at 10X of the final concentration, was added at a volume of 10 µl per well. After 2 h incubation, the plate was measured for baseline luminescence (Spectramax L, Molecular Devices) and 20 µl portions of 6X test compound, diluted in the assay buffer, were manually added. After an incubation for 3-5 minutes at room temperature, the plate was read for the second measurement. The second luminescence counts were normalized to the initial counts, and the fold-changes in the signals over the vehicle treatment were plotted for the G-protein dissociation response. Using the Prism 8 software (GraphPad Prism), the G-protein dissociation signals were fitted to a four-parameter sigmoidal concentration-response curve, from which the pEC_50_ values (negative logarithmic values of EC_50_ values) were used to calculate the mean and SEM. The pEC_50_ values for PACAP-17 were calculated by restraining the “Shared values for all datasets” for the “Top” and “Bottom” parameters, using both PACAP-17 and PACAP-38.

## Flow cytometry analysis

G_q/11_-deficient HEK293 cells^34^ were seeded in a 12-well culture plate at a concentration of 2 × 10^5^ cells ml^−1^ (1 ml per well), one day before transfection. The transfection solution was prepared by combining 2 µl of the polyethylenimine solution (1 mg ml^−1^) and 500 ng of a plasmid encoding the FLAG epitope-tagged GPCR in 100 µl of Opti-MEM. One day after transfection, the cells were collected by adding 100 μl of 0.53 mM EDTA-containing Dulbecco’s PBS (D-PBS), followed by 100 μl of 5 mM HEPES (pH 7.4)-containing Hank’s Balanced Salt Solution (HBSS). The cell suspension was transferred to a 96-well V-bottom plate and fluorescently labeled with an anti-FLAG epitope (DYKDDDDK) tag monoclonal antibody (Clone 1E6, FujiFilm Wako Pure Chemicals; 10 μg/ml diluted in 2% goat serum- and 2 mM EDTA-containing D-PBS (blocking buffer)) and a goat anti-mouse IgG secondary antibody conjugated with Alexa Fluor 488 (ThermoFisher Scientific, 10 μg/ml diluted in the blocking buffer). After washing with D-PBS, the cells were resuspended in 200 μl of 2 mM EDTA-containing-D-PBS and filtered through a 40-μm filter. The fluorescent intensity of single cells was quantified by an EC800 flow cytometer equipped with a 488 nm laser (Sony). The fluorescent signal derived from Alexa Fluor 488 was recorded in an FL1 channel, and the flow cytometry data were analyzed with the FlowJo software (FlowJo). Live cells were gated with a forward scatter (FS-Peak-Lin) cutoff at the 390 setting, with a gain value of 1.7. Values of mean fluorescence intensity (MFI) from approximately 20,000 cells per sample were used for analysis.

## Data Availability

The raw image of the PAC1R-miniGsβ_1_γ_2_-Nb35 complex after motion correction has been deposited in the Electron Microscopy Public Image Archive, under accession code XXXX. The cryo-EM density map and atomic coordinates for the PAC1R-mini-Gs-Nb35 complex have been deposited in the Electron Microscopy Data Bank and the PDB, under accession codes XXXX and ZZZZ, respectively.

---

**Supplementary Fig. 1.**
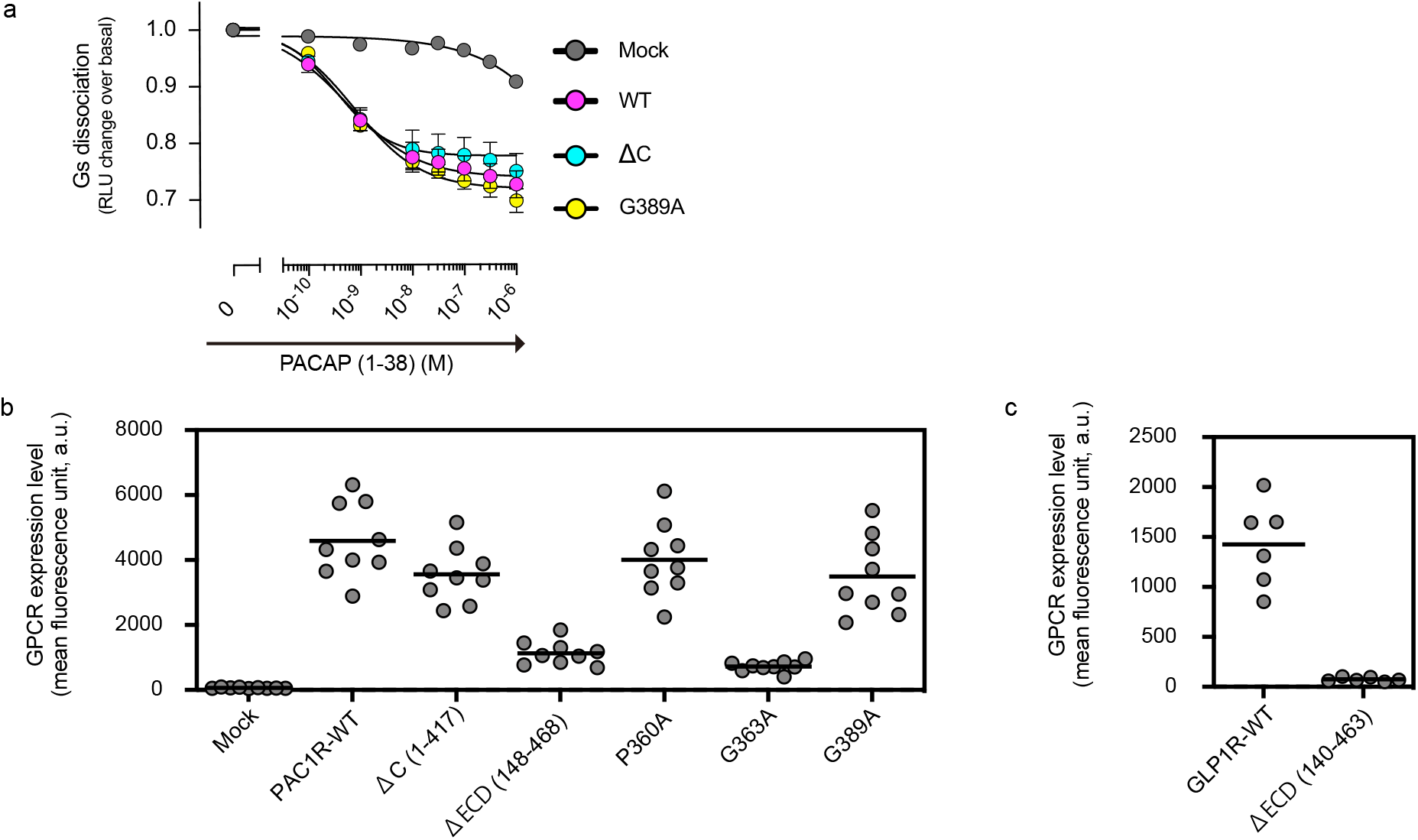
Functional characterization of mutant PAC1 receptors. **a**, PACAP-induced Gs activation, measured by the NanoBiT-G-protein dissociation assay. Cells transiently expressing the NanoBiT-Gs, along with the indicated PAC1R construct, were treated with PACAP (1-38) and the change in the luminescent signal was measured. **b, c**, Cell surface expression of the PAC1R and the GLP1R constructs. Cells transiently expressing the indicated FLAG epitope-tagged PAC1R constructs (**b**) or the GLP1R constructs (**c**) were labeled with an anti-FLAG tag antibody along with an Alexa488-conjugated secondary antibody, and the fluorescent signals from individual cells were measured by a flow cytometer.

**Supplementary Fig. 2.**
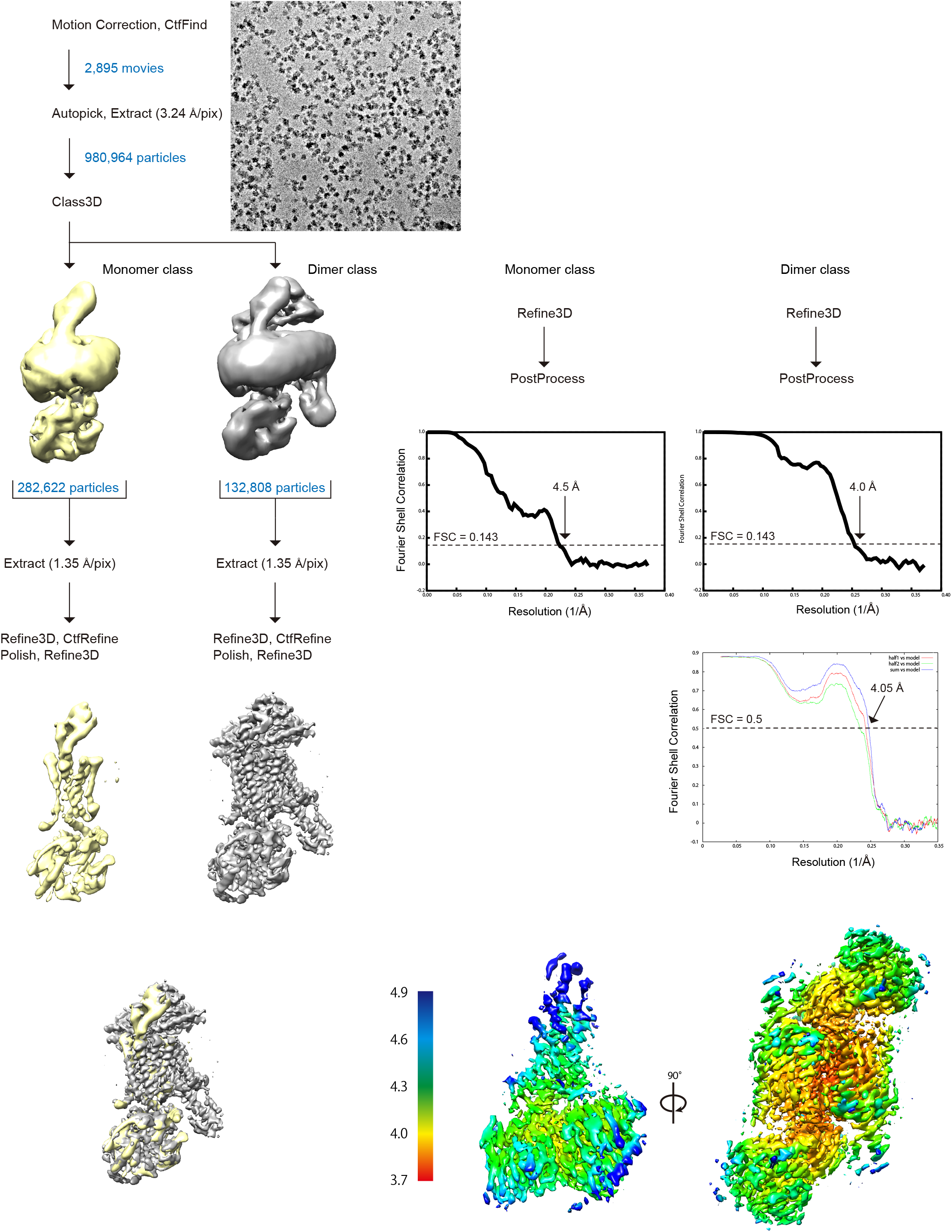
Cryo-EM analysis. Flow chart of the cryo-EM data processing for the PACAP–PAC1R–G_s_ complex, including particle projection selection, classification, and 3D density map reconstruction. Details are provided in the Methods section.

**Supplementary Fig. 3.**
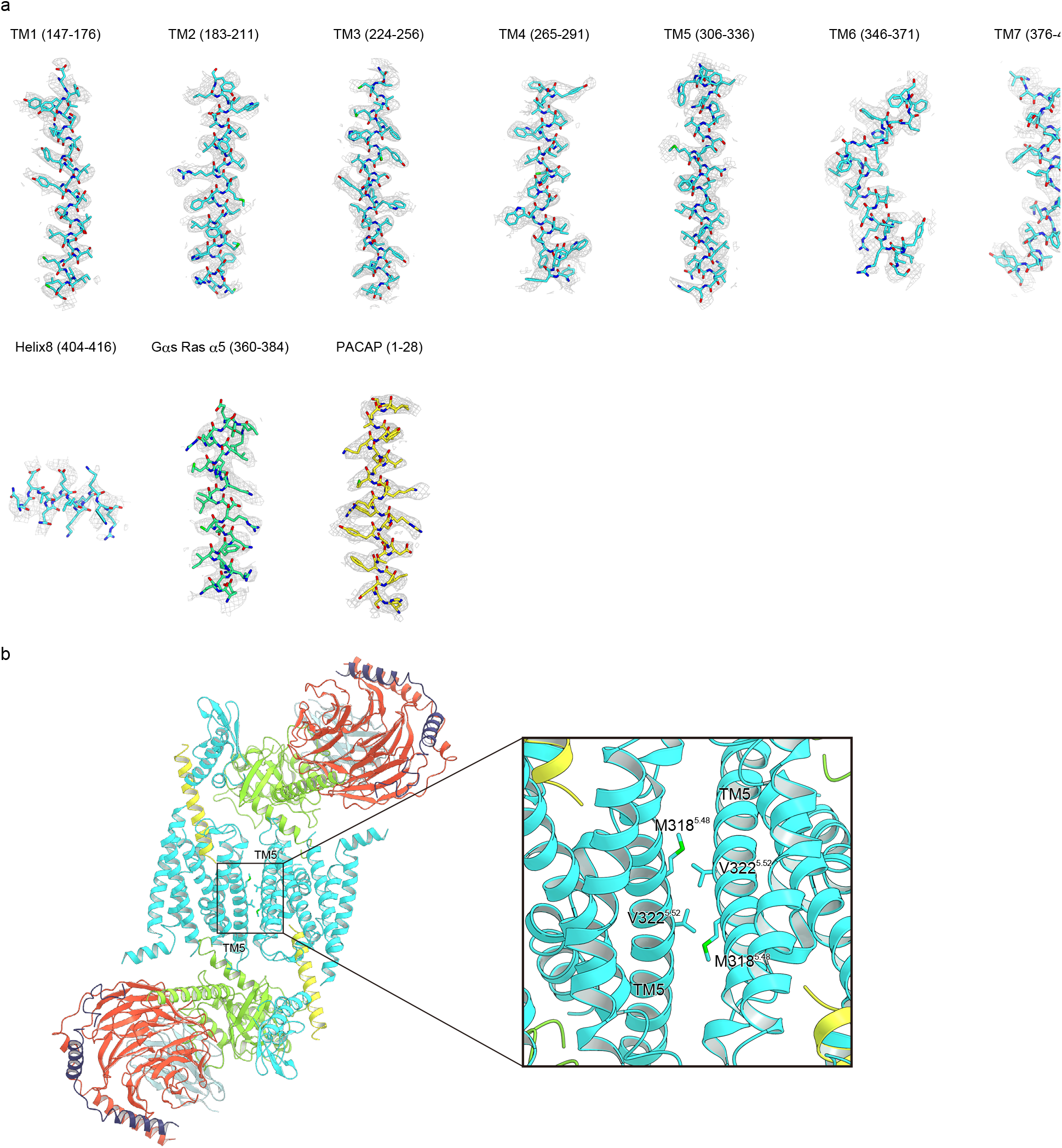
Map/model quality. **a**, The cryo-EM density map and model are shown for PACAP, including all seven transmembrane α-helices, ECD, and α5 of Gαs. **b**, Dimer interface. The complexes are shown as ribbon representations, colored as in Fig. 1a. The side chains of V318^5.48^ and M322^5.52^ are shown as sticks. Two complexes form an anti-parallel dimer with C2 symmetry in the detergent micelles. This dimer does not reflect the physiological condition, but is produced during the sample preparation. The molecular packing of the two complexes in the dimer class is mediated through only a weak hydrophobic contact between V318^5.48^ and M322^5.52^. Therefore, the dimerization minimally affects the conformation of the Gs-complexed PAC1R structure.

**Supplementary Fig. 4.**
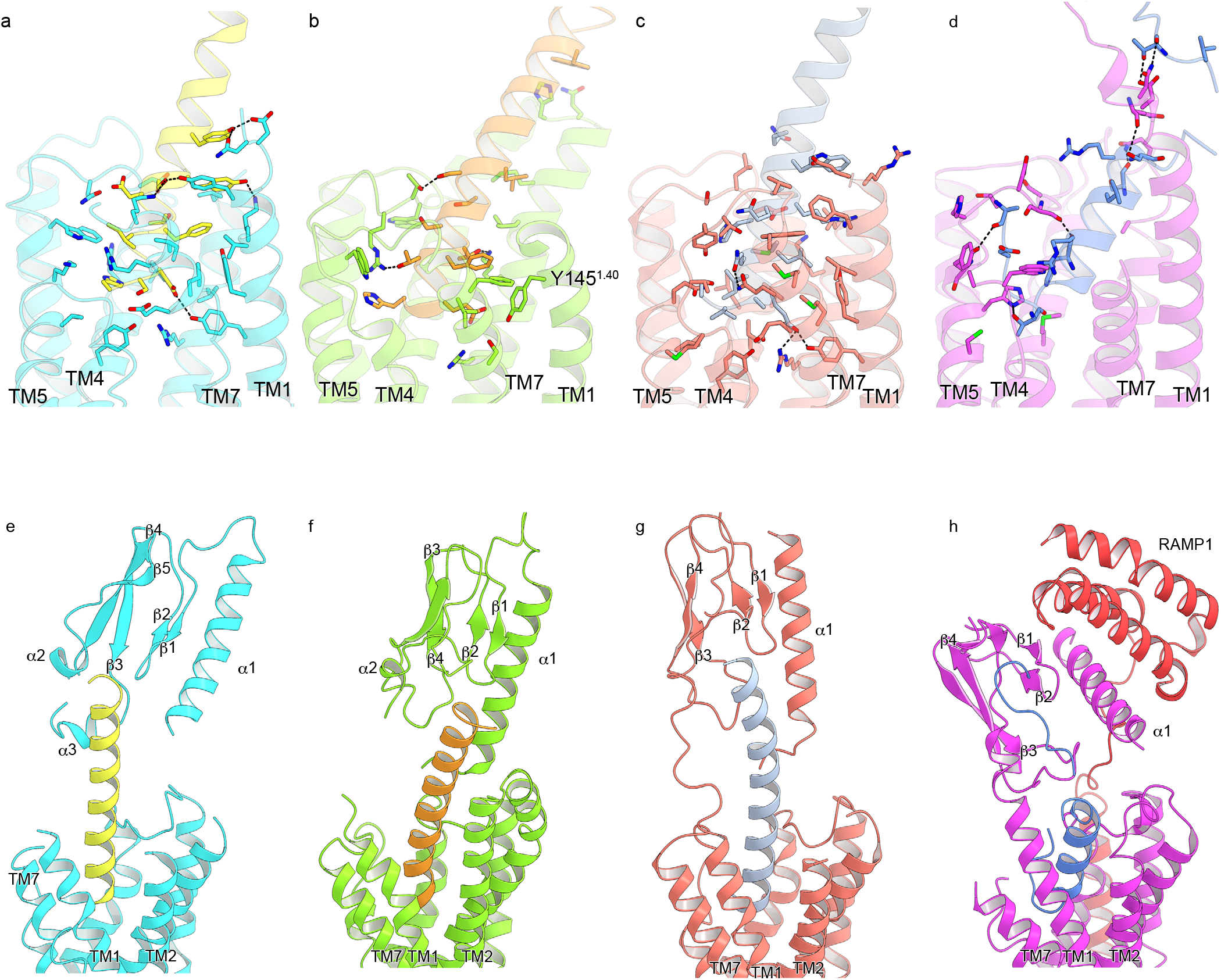
Comparison of peptide binding interactions in class B GPCRs. **a-d**, Ligand binding interactions with the TMDs in the class B GPCR structures (**a**, PAC1R, **b**, GLP1R, **c**, PTH1R, and **d**, CGRP). Hydrogen bonding interactions are indicated by black dashed lines. PACAP forms extensive hydrogen-bonding interactions with TM1, whereas GLP1 forms only a hydrophobic contact with Y145^1.40^. PTH also interacts with TM1; however, these interactions are mainly hydrophobic. **d-f**, Relative positions of the peptide ligands and the ECDs in the class B GPCR structures (**e**, PAC1R, **f**, GLP1R, **g**, PTH1R, and **h**, CGRP).

**Supplementary Fig. 5.**
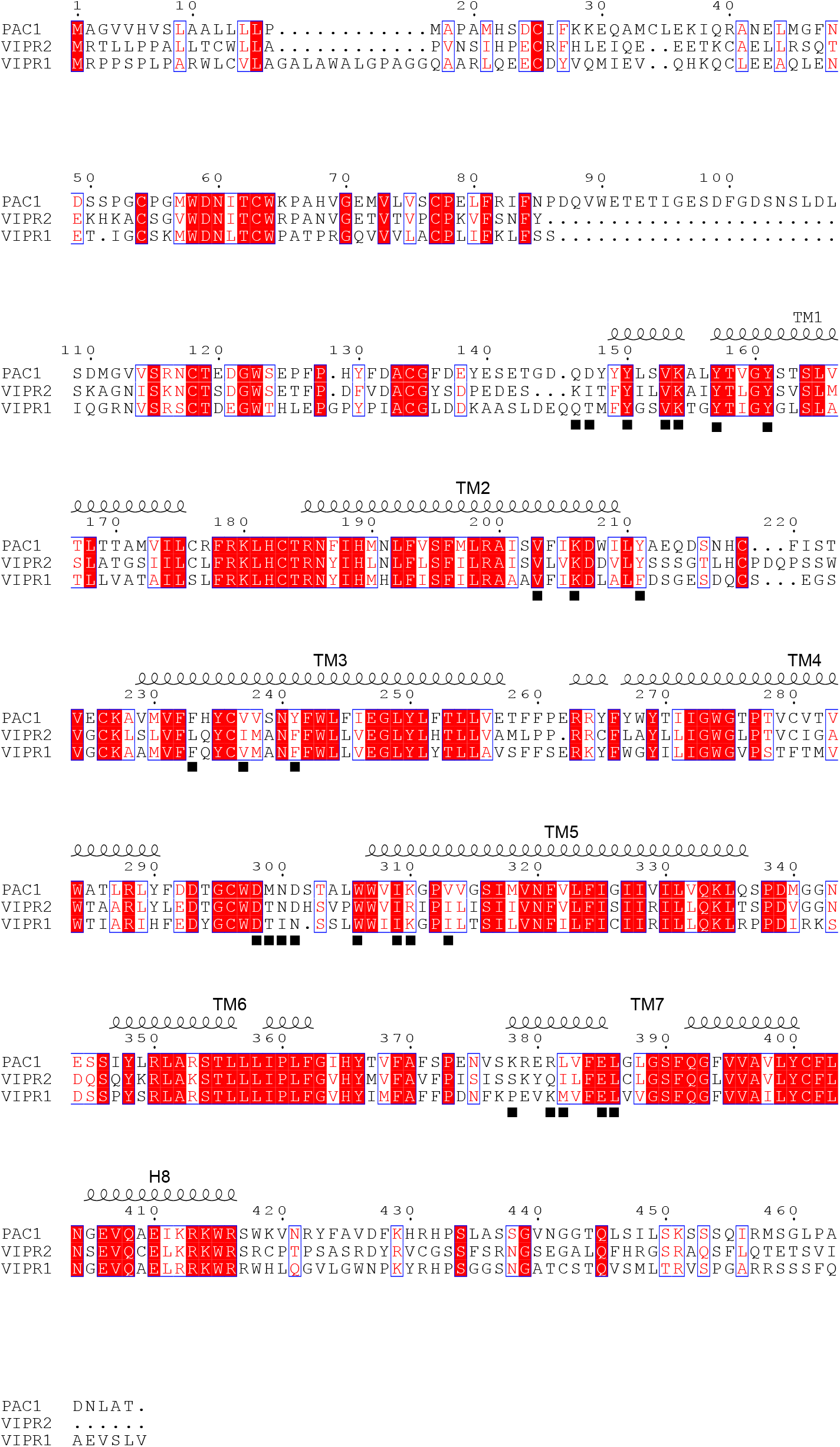
Sequence alignment of PAC1R, VPAC1R, and VPAC2R. Amino-acid sequences of the transmembrane domains of the human PAC1R (UniProt ID: P41586), VPAC1R (P32241), and VPAC2R (P25101). Secondary structure elements for α-helices and β-strands are indicated by cylinders and arrows, respectively. Conservation of the residues between the PACAP receptors is indicated as follows: red panels for completely conserved, red letters for partly conserved, and black letters for not conserved. The residues involved in the PACAP binding are indicated by squares.

**Supplementary Table 1.** Interactions of the PACAP N-terminal helix with the PAC1R-TMD. Residues within 4.0 Å are shown.

| PACAP | PAC1R | Interaction | Conserved |
| --- | --- | --- | --- |
| His1 | Val237 | Hydrophobic interaction |  |
|  | Tyr241 |  |  |
|  | Trp306 |  |  |
|  | Ile309 |  | C |
|  | Lys310 |  |  |
|  | Val313 |  |  |
| Ser2 | Glu385 | Hydrogen bond |  |
|  | Leu382 | Hydrophobic interaction |  |
|  | Leu386 |  | C |
| Asp3 | Tyr161 | Hydrogen bond | C |
|  | Val203 | Hydrophobic interaction | C |
|  | Phe233 |  |  |
|  | Leu386 |  | C |
| Gly4 | Asn300 | Hydrophobic interaction |  |
|  | Trp306 |  | C |
| Ile5 | Lys378 | Hydrophobic interaction |  |
|  | Arg381 |  |  |
|  | Leu382 |  |  |
| Phe6 | Tyr150 | Hydrophobic interaction | C |
|  | Val153 |  |  |
|  | Lys154 |  | C |
|  | Tyr157 |  | C |
|  | Leu382 |  |  |
|  | Leu386 |  | C |
| Thr7 | Lys206 | Hydrophobic interaction | C |
|  | Tyr211 |  |  |
|  | Asp298 |  | C |
| Asp8 | Asp298 | Hydrophobic interaction | C |
|  | Met299 |  |  |
|  | Asn300 |  |  |
| Ser9 | Tyr150 | Hydrogen bond | C |
|  | Lys378 |  |  |
| Tyr10 | Lys154 | Hydrogen bond | C |
|  | Tyr150 | Hydrophobic interaction | C |
|  | Tyr211 |  |  |
| Ser11 | Lys206 | Hydrophobic interaction | C |
|  | Tyr211 |  |  |
|  | Asp298 |  | C |
|  | Met299 |  |  |
| Arg12 | Asp301 | Electrostatic interaction |  |
| Tyr13 | Gln146 | Hydrophobic interaction |  |
|  | Asp147 | Hydrogen bond |  |
| Arg14 | Leu210 | Hydrophobic interaction |  |
|  | Tyr211 |  |  |
| Lys15 | Met299 | Hydrophobic interaction |  |
| Met17 | Asp147 | Hydrophobic interaction |  |

